# Long-term survival of chronically demyelinated axons in the *taiep* rat

**DOI:** 10.64898/2026.11.02.763294

**Authors:** Nathaniel Langan, Ben K. August, Katie J. Chapple, Hazel Allardyce, Dan Radecki, Ian D. Duncan, Julia M Edgar

## Abstract

Multiple sclerosis (MS) is an inflammatory demyelinating disease of the CNS in which neuroaxonal loss underlies permanent neurological disability. Axon loss in MS has been attributed to several causes, including loss of trophic/metabolic support from the myelin sheath. However, the contribution of chronic demyelination to axonal demise has been difficult to ascertain, due largely to the lack of long-lived models and uncertainty about the location of the primary injury site in long fibre tracts in MS patients. Here, we examine chronically demyelinated axons using light and electron microscopy of spinal cord tracts of 1- to 24-month-old *taiep* rats, in which the mutated gene, *Tubb4a* p.A302T, is expressed highly in oligodendrocytes. We find that small diameter axons in the dorsal column can survive for many months following demyelination. This is associated with significantly increased densities of oligodendroglia, and supernumerary glial cell processes. We hypothesise that trophic and metabolic support, sufficient to support axon survival, can be achieved through close contact between axons and glia. While efforts to restore myelin in MS remain crucial to recovering normal neurological function, myelin *per se* may not be a prerequisite for axonal survival under all circumstances.

## Introduction

MS is a chronic inflammatory demyelinating disease of the CNS in which neuro-axonal loss is generally considered the most important correlate of persistent disability (Ferguson et al., 1997; Trapp et al., 1998; De Stefano et al., 1999; Tallantyre et al., 2010; Franklin et al., 2012; Singh et al., 2017; Schirmer et al., 2019; Zoupi et al., 2021; Kuhlmann et al., 2023) and is reflected in considerable brain atrophy (De Stefano et al., 2010). Axonal injury in MS is widespread, occurring at focal sites of active demyelination, but also in neighbouring normal appearing white matter (NAWM), and in inactive demyelinated plaques. In general, axonal injury and loss can result from a primary insult at the level of the axon or the neuronal soma, leading to Wallerian degeneration distal to the injury, or following retraction of the axon terminus, leading to a dying back axonopathy (Gillingwater and Ribchester, 2001; Conforti et al., 2007; Brady and Morfini, 2010; Singh et al., 2017).

In MS white matter, axonal injury is generally understood to initiate at the level of white matter lesions (Ferguson et al., 1997; Trapp et al., 1998; Evangelou et al., 2000; Kuhlmann et al., 2002; Tallantyre et al., 2009; Singh et al., 2017) where, reports indicate, there is greater axon loss than in neighbouring NAWM (Tallantyre et al., 2009); albeit not all studies agree (Lovas et al., 2000). Evidence from patient studies and experimental models of myelin-associated disease suggest that in active and chronic active lesions, axonal injury is mediated by lipid-soluble inflammatory factors produced by macrophages and/or by granzyme and perforin produced by cytotoxic T cells (Smith et al., 2001; Kuhlmann et al., 2002; Edgar et al., 2010; Haider et al., 2011; Nikić et al., 2011; Sorbara et al., 2014; Schäffner et al., 2021; Green et al., 2025). Importantly, several of these studies show that axonal injury can occur independently of myelin loss (Smith et al., 2001) (Bitsch et al., 2000; Edgar et al., 2010; Nikić et al., 2011; Sorbara et al., 2014; Schäffner et al., 2021). In inactive, chronically demyelinated lesions, axon demise has been attributed to a state of “virtual hypoxia” brought about by energy insufficiency and increased energy demand (Trapp and Stys, 2009) and/or a loss of trophic support from the myelinating cell (Nave and Trapp, 2008; Trapp and Nave, 2008; Trapp and Stys, 2009; Mahad et al., 2015; Duncan et al., 2021).

However, the relationship between chronic demyelination *per se*, and axonal demise (Lovas et al., 2000; Bjartmar and Trapp, 2003) remains challenging to determine, and in MS, an ongoing role for inflammation is difficult to exclude (Lassmann, 2003). The paucity of experimental studies examining the relationship between chronic demyelination and axon demise stems from the rarity of long-lived models of chronic demyelination, and the efficiency of remyelination in many spontaneous and induced demyelinating models (Franklin and Ffrench-Constant, 2017).

Previously, using a demyelinating mouse model of Pelizaeus Merzbacher disease (PMD) due to proteolipid protein gene (*Plp1)* overexpression, we showed that focal axonal injury was more prevalent in macrophage-rich regions undergoing active demyelination, compared to regions of the same axons that were already fully a-or de-myelinated (Edgar et al., 2010). Furthermore, quantification of axons from transmission electron microscopy (TEM) micrographs, demonstrated that axon numbers were not reduced compared to WT, indicating that axons can survive acute demyelination; many could conduct electrical impulses, albeit slowly (Edgar et al., 2010). However, the short (4-months) lifespan of this model prevented longer-term tracking of the fate of these demyelinated axons. More recently, in a cell type specific myelin regulatory factor (*Myrf*) knockout mouse, Duncan et al., showed by TEM, that 20 weeks following genetically induced demyelination, axon numbers were unaltered despite that ∼20% remained demyelinated (Duncan et al., 2024). In contrast, ∼60% and 45% decreases, respectively, were reported in optic nerve axons in two independent long-lived demyelinating models; 12-months post-demyelination in the *taiep* rat (Wilkins et al., 2010) and at 9 months post-toxin-induced demyelination in macaque (Sarrazin et al., 2022). Notably however, in these latter reports, axons were identified by neurofilament immunohistochemistry.

Together, these optic nerve data suggest that the fate of demyelinated axons is, at least in part, determined by the chronicity of the myelin loss. Notwithstanding, it remains possible that neurofilament immunostaining combined with light microscopy, as used in the latter two studies, fails to identify all surviving demyelinated axons.

Here we used TEM to study the fate of chronically demyelinated spinal cord axons in the long-lived *taiep* rat. *Taiep* harbours a single point mutation in the tubulin class IVa (*Tubb4a*) gene, which encodes the microtubule subunit protein β-Tubulin 4A (Duncan et al., 2017). In the CNS, *Tubb4a* is most highly expressed in oligodendrocytes (Terada et al., 2005; Wu et al., 2009), where it is localised to cell processes and cytosolic spaces of myelin (Terada et al., 2005; Edgar et al., 2021; Chapple et al., 2026), which provide a transport route across myelin (TRAM; (Chapple et al., 2026)). Knocking out *Tubb4a* specifically in oligodendrocytes has no overt consequences for optic nerve myelin volume or structure (Chapple et al., 2026), suggesting the *taiep* mutation causes a toxic gain of function.

Consistent with this, *taiep* oligodendrocytes accumulate microtubules *in vitro* and *in vivo*, disrupting intracellular trafficking of myelin basic protein (*Mbp*) mRNA (Song et al., 1999; O’Connor et al., 2000) and various myelin proteins (Möller et al., 1997; Song et al., 1999). These changes are associated with hypomyelination and progressive demyelination of CNS white matter tracts, most notably those tracts that comprise small diameter axons (Duncan et al., 1992, 2017; Lunn et al., 1997).

Here we demonstrated that, in 24-month-old *taiep* rats, axonal densities remained comparable to WT, despite extensive demyelination of small calibre axons in the cervical spinal cord. The surviving axons were interspersed with glial processes and increased densities of oligodendroglia. Our data suggest that compact myelin *per se* is not required for the survival of (previously) myelinated axons when alternative routes for trophic and metabolic support are available.

## Methods and Materials

### Animals

Animal experiments comply with the current laws of the USA and the UK, in which they were performed.

Animal experimentation was approved by the University of Madison - Wisconsin Animal Care and Use Committee (Madison, USA). Sprague-Dawley wild -type (WT) and *taiep* mutant rats were bred and maintained at the University of Madison Charmany Animal Facility with *ad libitum* access to food and water. To avoid triggering seizure activity, affected rats were maintained in a quiet environment with minimal disturbance. Rats were euthanised by i.v. injection of 400mg/ml pentobarbital (BioVeta, Cat. No. 237) at a dose of 1ml/kg body weight. Three WT and three *taiep* rats were euthanised at each of the following ages: 1, 3, 6, 9 and 24 months. Affected and WT rats were identified in litters born from heterozygous *taiep* rat matings by their clear phenotype (*taiep*) or PCR (WT) (Duncan et al., 2017).

Adult Sprague-Dawley rats were also maintained at the University of Glasgow Biological Services facility, under project licence PPL P78DD6, and experiments approved by the local ethical review committee. Rats were housed in standard plastic cages. They were maintained with a 12-h dark-light cycle at 19-23°C and provided food and water *ad libitum*.

### Tissue preparation for electron microscopy

Euthanised rats were fixed by perfusion with physiological saline solution (0.154 M NaCl) followed by glutaraldehyde-paraformaldehyde solution (5% glutaraldehyde, 4% paraformaldehyde [PFA], 9 mM CaCl2, 80 mM sodium cacodylate) as previously described (Edgar et al., 2020). Whole spinal cord was dissected and postfixed for 24 hours at 4°C. The cervical spinal tissue was cut into 2 mm cubed blocks before secondary fixing in 1% osmium tetroxide in cacodylate buffer (4 mM OsO4, 0.07 M sodium cacodylate, 0.07 M NaCl, 0.0005 M CaCl2, 0.001 M MgCl2). Tissue blocks were then embedded in araldite resin (Agar Scientific, cat. no. AGR1030). Transverse sections were cut at 1 μm for light microscopy or 50 nm using a Leica UC6 ultramicrotome with a 45° 4 mm edge diamond knife (DiATOME, cat. no. DU4540). One section was cut from the dorsal column of each animal, and included the fasciculus gracilis (F. gracilis), fasciculus cuneatus (F. cuneatus) and corticospinal tracts (CST). One-micron sections were stained with toluidine blue. Only one F. cuneatus sample was obtained successfully from both WT and *taiep* 1-month-olds due to tract underdevelopment, so this condition was excluded from quantitative analysis.

### Transmission Electron Microscopy (TEM)

50 nm thick tissue sections were placed on 200 square formvar-coated copper TEM grids (Sigma-Aldrich, cat. no. 930318) and stained with Reynold’s lead citrate (0.1 M lead citrate, 0.2 M sodium citrate). Sections were visualised using a JEM-1400Flash transmission electron microscope (JEOL., Tokyo, Japan) set at 80 kV accelerating voltage. For quantification, micrographs were captured at 4400 X or 5600 X magnification using an inbuilt 2000x2000 CCD digital camera and saved as .tif files using software version 1.7.26.3016 (JEOL, Tokyo, Japan).

### Axon and mitochondrial quantification

To quantify myelinated and non-myelinated axons, rat spinal micrographs were imported as .tif files into Fiji (ImageJ2, version 2.14.0/1.54f) (Schindelin et al., 2012). A fixed region of interest (ROI) of defined size was applied to each image, and axons were quantified within and on south and west borders of the ROI and ascribed to myelinated or non-myelinated in the Fiji Cell Counter plugin. Three to eight micrographs per tract per animal, were quantified. Axons were identified as pale, approximately circular objects, 0.2-20 μm in diameter with microtubules and neurofilaments, to distinguish them from glial cell processes.

Axoplasm and axonal mitochondrial volumes were assessed using the ‘Grid’ tool in Fiji (Schindelin et al., 2012). Grid squares 2 μm^2^ were placed over each of 3-8 micrographs per tract, and intersections that occurred on axoplasm or axonal mitochondria were quantified in the Cell Counter plugin in Fiji. Mitochondria were identified based on having a double membrane, being approximately circular in shape and containing cristae.

Data collation was performed in Microsoft Excel. Myelinated and unmyelinated axon density was calculated for each tract per animal using each animal’s summed axon count and summed ROI area (total ROI) from the 3-8 micrographs per tract. Axonal density per mm^2^ was calculated as follows:

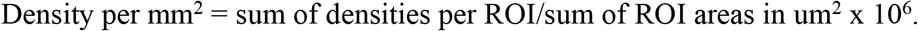

Mitochondrial volume was calculated by expressing the mitochondrial hit count as a percentage of axoplasmic + mitochondrial hit count.

Axon density, myelinated axon percentage and mitochondrial volume mean values were calculated from the three animals per age and genotype.

### Glial cell counts based on ultrastructural morphology

The total number of oligodendrocytes, astrocytes and microglial cells in the F. gracilis of three 24-month-old *taiep* and WT rats was determined on TEM evaluation. Identifying and classifying the three cell types was performed using standard ultrastructural criteria (Peters et al.) and each of the three cell types were easily distinguished. In addition, oligodendrocytes in the 24-month-old *taiep* rats had accumulation of microtubules which confirmed their identity. Cells were counted in three adjacent TEM grid squares in the centre of the F. gracilis in each rat.

### Immunohistochemistry of WT rat spinal cord

Rats were killed by intraperitoneal injection with Dolethal followed by exsanguination. The cervical spinal cord and associated vertebrae were isolated from the head and thorax and placed directly in 4% PFA (overnight incubation at 4°C). The cords were dissected the following day and incubated again in 4% PFA at 4°C overnight. The cords were then placed in 20% sucrose in PBS until they sank. The tissue was then frozen in liquid nitrogen-chilled isopentane in OCT (Sakura Finitek). Transverse sections were serially cut at 10 μm thickness on a cryostat (Epredia NX50 Rotary Cryotome) and collected 40 μm apart on Plus-charged slides (Thermo Fisher Scientific) then stored in a sealed slide box at -80°C.

### Immunohistochemistry

Sections were washed in PBS to remove the OCT then permeabilised using methanol (-20°C, 20 minutes). They were incubated for 1 h in blocking buffer (10% goat serum, 1% BSA, 0.1% Tween and 1.8.% sodium chloride in phosphate buffer). Sections were immunolabelled with rabbit anti-ASPA (1:1000; kindly gifted by the late James Garbern) and mouse anti-β-tubulin 4 (1:400; Abcam; ab11315) by incubation at 4°C overnight in blocking buffer. Bound primary antibodies were detected using Alexa 488 goat anti-mouse IgG1 and Alexa 568 goat anti-rabbit IgG (1:1000; Invitrogen) in blocking buffer for 1h Sections were mounted in Mowiol with DAPI (5 μg/mL).

Immunostaining for microglia was carried out as previously described (Zhang et al., 2003). Briefly, free-floating spinal cord sections from one to thirteen-months old animals were immunolabelled with an antibody to OX-42 (anti-CD11b; mouse IgG; Sero Labs).

### Counts of immunohistochemically stained oligodendrocytes

Non-overlapping images were captured across the entire dorsal columns at 40x magnification using an inverted Zeiss Observer Z1 widefield epifluorescence microscope equipped with an AxioCam MRc3, x0.63 Camera Adaptor, ZEN 2012 blue edition software using Zeiss plan-apochromat x40 objective in the red (β−tubulin 4), green (ASPA) and blue (DAPI) channels. Images were quantified using the Cell Counter plugin in Fiji. The blue and green channels were overlaid in Fiji and ASPA positive structures with a nucleus, were quantified. The red channel was then overlaid on the green and blue channels, and by switching channels off and on, it was straightforward to identify ASPA positive cells also positive for β−tubulin 4. Quantification was undertaken by two independent experimenters, on two cords, three sections (40 μm apart) per cord, and the experimenters’ values were averaged. ASPA was observed in cell bodies and nuclei and in some primary processes; β−tubulin 4 was observed in cell bodies and primary processes, but not in cell nuclei. β−Tubulin 4 was also seen in structures assumed to be myelin sheaths.

### Statistical testing

Statistical testing was performed in Prism (version 10.0.0., GraphPad Software Inc.). Axonal densities and axoplasm/mitochondrial volumes were analysed using a 2-way ANOVA followed by Šídák’s multiple comparisons test. Percent myelinated axons were analysed using multiple unpaired t-tests followed by a two-stage false discovery rate test (Benjamini, Krieger, and Yekutieli). A p value <0.05 was considered statistically significant. Statistical significance is indicated in Figs as: * *= p < 0.05, ** **= p < 0.01, \*\*\**** = < 0.001 or ‘****’ = p < 0.0001.

## Results

### *Taiep* rats exhibit a progressive neurological phenotype

*Taiep* rats exhibit tremor, ataxia and hind-limb paresis (Lunn et al., 1997). Neurological abnormality begins at around 30 days of age, with development of a tail tremor which progresses over the next month to involve the whole body. This tremor worsens and hind leg ataxia develops. By 6-9 months, the fine tremor is replaced by a coarse, whole-body tremor. Between 9-12 months, seizures begin, which at rest are infrequent, but handling or disturbance frequently initiates a seizure event. Here, with careful husbandry, we were able to maintain *taiep* rats until >24 months of age at which time these neurological signs persisted, and the animals were killed humanely according to local regulations.

### Progressive demyelination of small diameter dorsal column axons in *taiep*

As reported previously, *taiep* is characterised by early hypomyelination followed by a progressive loss of myelin in brain and spinal cord (Lunn et al., 1997; Duncan et al., 2017). At 24 months of age, in the cervical spinal cord, the dorsal columns appeared markedly more affected than the ventral and lateral columns, **Supplementary Fig. 1,** therefore we focussed our studies on the first of these. Using toluidine blue staining of transverse sections of the *taiep* cervical spinal cord dorsal columns at various ages, we confirmed our earlier observations demonstrating a progressive loss of myelin that is most marked in the corticospinal tract (CST) and fasciculus gracilis (F. gracilis) compared to the fasciculus cuneatus (F. cuneatus) (**Fig. 1a-b**). Strikingly, this remained so at 24 months (**Fig. 1c**). The CST and F. gracilis comprise small diameter descending and ascending fibres, respectively. Furthermore, the cervical F. gracilis contains the distal ends of the longest axons in the rodent CNS, which arise in the dorsal root ganglia (DRG) that serve the lower limbs and lower trunk. In contrast, the F. cuneatus contains a mixture of small, medium and larger diameter axons. These arise in DRGs serving the upper limbs and trunk and ascend through the upper spinal cord.

**Fig. 1.**
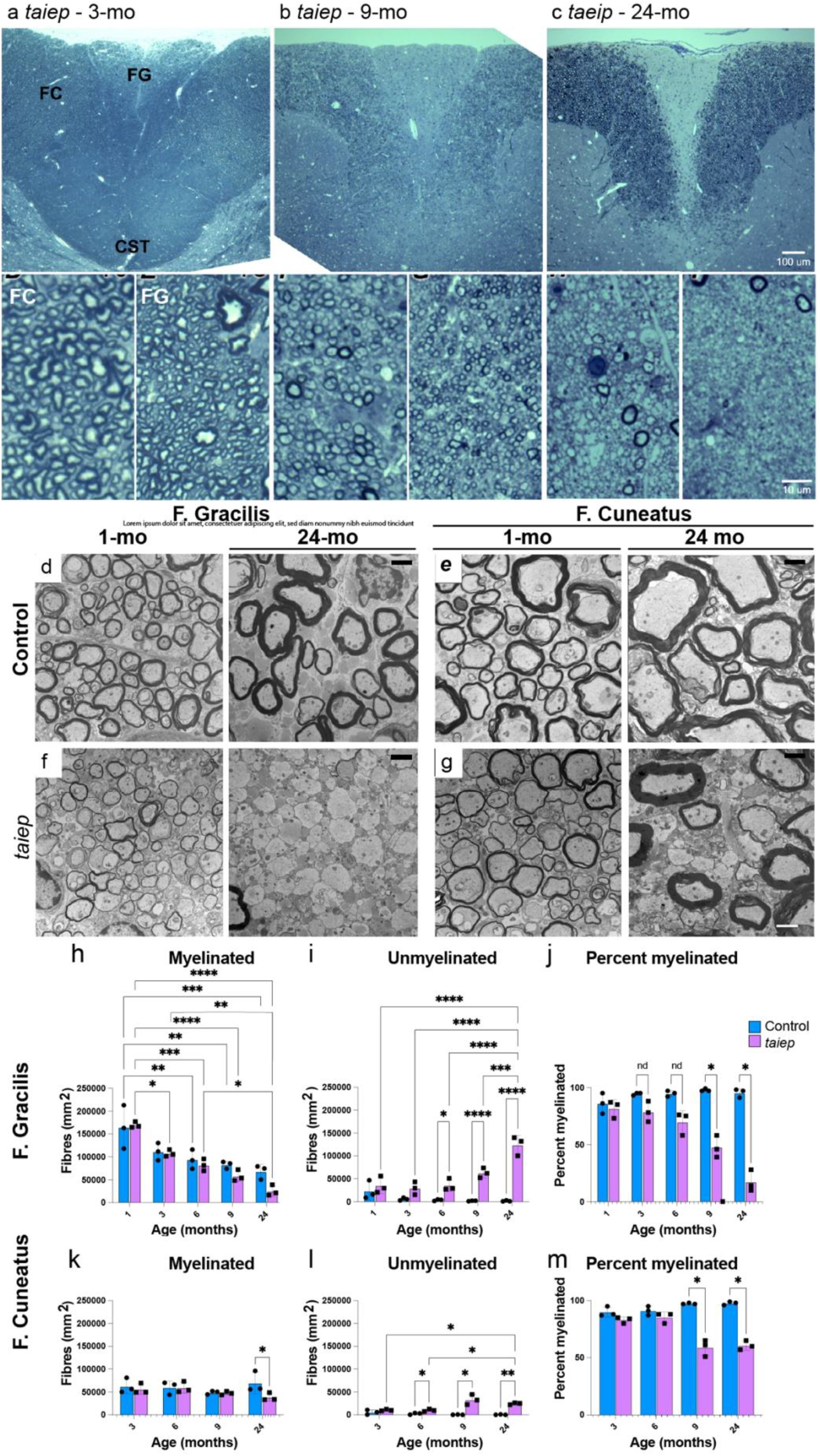
Progressive demyelination in *taiep* cervical cord dorsal columns a-c. Overviews (top) and higher magnification images (beneath) of toluidine blue stained resin sections of *taiep* cervical spinal cord dorsal column at 3-, 9- and 24 months. At 3 months of age, all three tracts appeared well myelinated. By 9 months there was relative pallor of the F. gracilis and CST. At 24 months, pallor was marked in the F. gracilis and CST, while the F. cuneatus retained many myelin sheaths. F. gracilis (FG), F. cuneatus (FC) and corticospinal tract (CST). **d&e.** TEM micrographs of WT F. gracilis and F. cuneatus at 1- and 24 months of age. **f&g.** TEM micrographs of *taiep* F. gracilis and F. cuneatus at 1- and 24 months of age. In *taiep*, the majority of F. gracilis axons was devoid of myelin whereas many F. cuneatus axons remained myelinated. **h.** In the F. gracilis, the densities of myelinated axons decreased significantly between 1 month and 6, 9 or 24 months in both groups, and between 1 and 3 months in *taiep*. In *taiep* there was also a significant decrease in densities between 3 or 6 months and 24 months. **i.** In the *taiep* rat F. gracilis, there was a significant increase in the densities of non-myelinated axons between 1-, 3-, 6- or 9 months and 24 months and between WT and *taiep* at 6-, 9- and 24 months of age. **j.** The percentage of axons that were myelinated was significantly reduced in *taiep* compared to WT at 9- and 24 months. **k.** In the F. cuneatus, the densities of myelinated axons is similar in *taiep* and WT at 3-, 6- and 9 months, but significantly reduced in *taiep* at 24 months. **l.** In the F. cuneatus, there was a significant increase in the densities of non-myelinated axons between 3 or 6 months and 24 months in *taiep* and in *taiep* compared to WT at 6-, 9- and 24 months. **m.** The percentage of axons that were myelinated was significantly reduced in *taiep* rat compared to WT at 9- and 24 months. All graphs: Densities were analysed using a 2-way ANOVA followed by Šídák’s multiple comparisons test. The percentage myelinated axons was analysed using multiple unpaired t-tests followed by a two-stage false discovery rate test (Benjamini, Krieger, and Yekutieli). Each black dot represents a single animal, bars represent mean +/-SD.

To compare myelin status in diverse tracts over time (**Fig. 1d-g**), we quantified densities of myelinated and non-myelinated axons in TEM micrographs of the cervical F. gracilis and F. cuneatus in *taiep* and WT rats between 1 or 3, respectively, and 24 months of age (**Fig. 1h-m)**.

F. gracilis: Comparing WTs and *taiep* to themselves at increasing ages, we found that the densities of myelinated axons decreased significantly in both groups between 1-month and 6-, 9-or 24 months. This is most likely due to increasing fibre diameter as the animals reach adulthood. In *taiep*, there was also a significant decrease at 3 months compared to 1-month and at 24 months compared to 3-or 6 months (**Fig. 1h**). Simultaneously, the densities of unmyelinated axons increased significantly in *taiep* at 24 months compared to 1-, 3-, 6-or 9 months (**Fig. 1i**). This suggests the reduction of myelinated axons is due not to axon loss, but to demyelination. Comparing WT and *taiep* to each other, we observed significantly more demyelinated axons in *taiep* at 6-, 9- and 24 months (**Fig. 1i**) Thus, in *taiep*, myelinated axons as a percentage of all axons tended to decrease from 3 months, with differences compared to WT reaching statistical significance at 9- and 24 months (**Fig. 1j**).

F. cuneatus: Comparing WT and *taiep* to themselves over time from 3 months, we found no differences in the densities of myelinated axons with increasing age (**Fig. 1k**). However, the densities of non-myelinated axons increased significantly in *taiep* at 24 months compared to 3- or 6 months (**Fig. 1l**). Comparing WT and *taiep* to each other, myelinated axon densities were significantly reduced in *taiep* compared to WT at 24 months (**Fig. 1k**) while densities of unmyelinated axons were slightly but significantly increased compared to WT at 6-, 9- and 24 months (**Fig. 1l**). These changes resulted in a significant reduction in the percentage of myelinated axons at 9- and 24 months in *taiep* compared to WT (**Fig. 1m**).

### Profound loss of myelin in the *taiep* rat CST

In contrast to the F. gracilis and the F. cuneatus, in which ∼10 and ∼60 % respectively, of axons remained myelinated at 24 months, virtually all CST axons were devoid of a myelin sheath at 24 months (**Fig. 2**). These axons were tightly packed with no evidence of axon loss; occasional astrocyte processes were noted and a few scattered axons had myelin sheaths (**Fig. 2d**). Notwithstanding, a substantial proportion of the predominantly small diameter axons in the taiep CST were myelinated at 3 months (**Fig. 2c**), indicating that the absence of myelin at 24 months was due primarily to loss of pre-existing myelin.

**Fig. 2.**
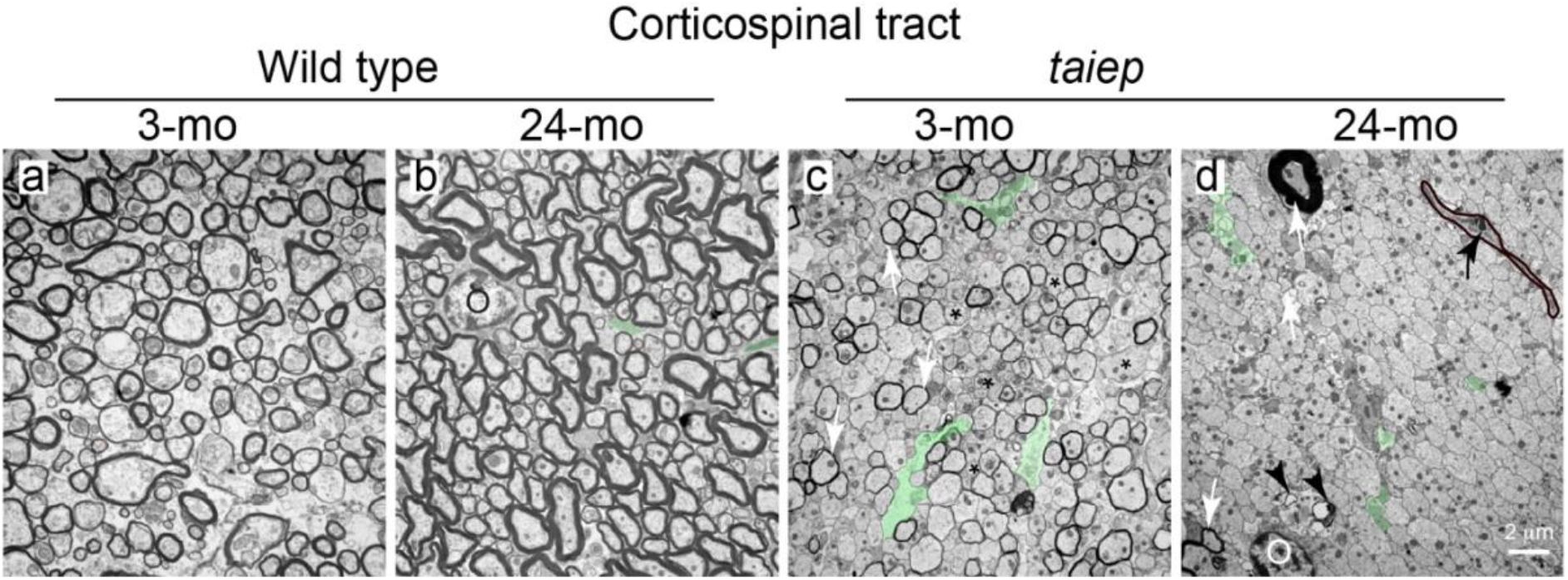
The *taiep* rat corticospinal tract is almost completely devoid of myelin at 24 months a&b. In WT, the neuropil was tightly packed with small diameter myelinated axons in which the myelin sheaths thickened with time from 3 to 24 months. O: oligodendrocyte. **c.** In contrast, in the 3-month-old *taiep* rat, only a proportion of axons were myelinated (arrows), and single or small groups of unmyelinated axons were present (asterisks). **d.** By 24 months there was a striking loss of myelin with only rare axons retaining a sheath (white arrows). Notably, axons otherwise appeared normal. In the upper left of the image, an axon in surrounded by an abnormally thick sheath. Small amounts of myelin debris remained (black arrowheads). A microglial process containing a cholesterol crystal (black arrow) is delineated in black in the upper right of the image.

### Differences in fibre susceptibility to demyelination are unlikely due to heterogeneity in oligodendrocyte expression of β−tubulin 4

As we reported previously (Lunn et al., 1997), small diameter *taiep* axons appeared particularly susceptible to demyelination. This was especially obvious at 24 months at the border between the F. gracilis (principally small diameter axons) and the F. cuneatus (mixture of small and larger diameter axons) (**Fig. 3a**), and in the F. cuneatus itself where large diameter axons remained myelinated at 24 months, but small diameter axons were demyelinated or surrounded by an abnormally thin sheath (**Fig. 3b**). While the morphological diversity of oligodendrocytes was first recorded in the early 20th century by Pío del Río-Hortega, the emergence of single-cell and single-nucleus RNA sequencing technologies revealed that also at the level of the transcriptome, mature oligodendrocytes represent a heterogeneous population spanning at least 6 distinct transcriptional states (Zeisel et al., 2015; Marques et al., 2016). This raises the possibility that the unaffected large diameter fibres in *taiep* F. cuneatus, are myelinated by oligodendrocytes lacking (‘mutant’) β-tubulin 4. To address this, we immunolabelled adult WT rat spinal cord with anti-aspartoacylase (ASPA), a marker of mature oligodendrocytes (Madhavarao et al., 2004) (Klugmann et al., 2003) and anti-β-tubulin 4. We found that across the entire dorsal columns, 93.7 % (+/-4.0 SD; n = 2) of ASPA +ve cells were also β-tubulin 4 +ve (**Fig. 3b-e**). In contrast to our hypothesis, the few ASPA +ve somata that lacked or only weakly expressed β-tubulin 4, were located in the CST, not the F. cuneatus.

**Fig. 3.**
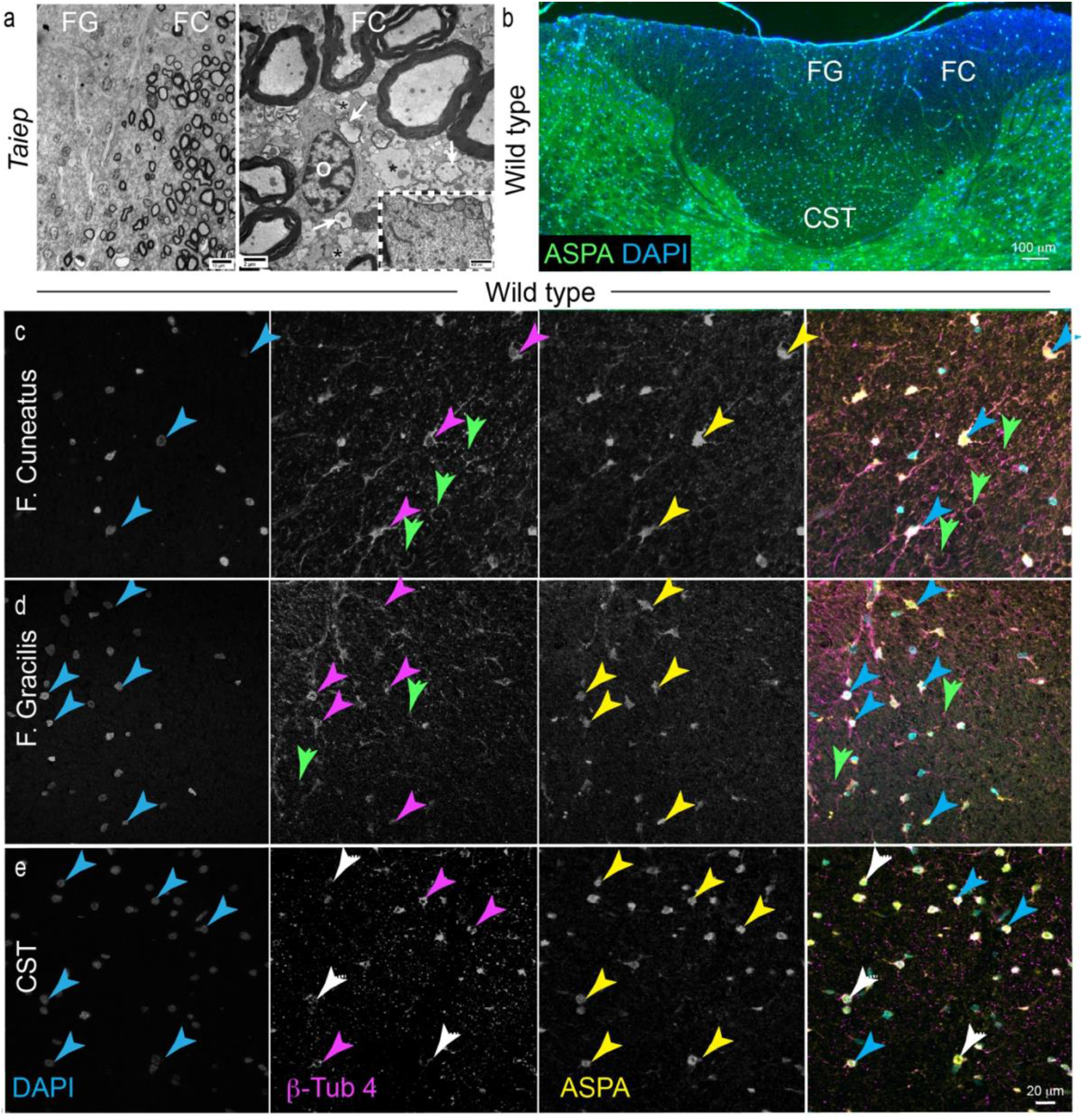
Myelin preservation on large diameter fibres in *taiep* rats appears unrelated to β−tubulin 4 expression in oligodendrocyte somata a. Left: TEM micrograph of the F. cuneatus/F. gracilis interface. There is a clear demarcation between the two tracts, with the larger axons of the F. cuneatus being normally myelinated in contrast to the complete demyelination of the small axons in the F. gracilis. Scale bar: 10 μm. Right: TEM micrographs illustrating the difference in myelination between large and small diameter axons in the F. cuneatus of old *taiep* rats. In contrast to the F. gracilis which has only small diameter axons, the large axons of the F. cuneatus have normal thickness myelin sheaths. However, small axons are either demyelinated (asterisk) or hypomyelinated (arrows). These axons are adjacent to an oligodendrocyte (O) that has accumulated microtubules and microtubule/smooth endoplasmic reticulum arrays (inset). F. gracilis (FG), F. cuneatus (FC) and corticospinal tract (CST). Scale bars: 2 μm and 400 nm. **b.** Immunohistochemical staining of dorsal columns and adjacent grey matter of an adult WT rat showing cell nuclei (DAPI) and ASPA +ve oligodendrocyte somata (green); the latter are seen in both white and grey matter and are particularly densely packed in the CST. **c&d**. In the F. cuneatus and F. gracilis, all ASPA +ve oligodendrocyte somata were also positive for β−tubulin 4 (examples indicated by arrowheads in the three colour channels and the overlay). Occasionally, β−tubulin 4 staining formed small circular structures that lacked nuclei (green arrows in magenta channel) suggesting these are paranodal loops where myelin- located microtubules often encircle the axon (Edgar et al., 2021; Chapple et al., 2026). **e.** In the CST, occasional ASPA +ve somata lacked β−tubulin 4 staining or the signal was particularly faint (white arrows in magenta channel and in overlay). However, the majority of ASPA +ve oligodendrocyte somata were also positive for β−tubulin 4 (examples indicated by arrowheads in the three colour channels and the overlay). Scale bar c-e: 20 μm. Two rats examined.

These data support the suggestion that β-tubulin 4 expression *per se* does not render oligodendrocytes vulnerable to the *taiep* mutation and support our previous suggestion that microtubule-dependent trafficking of components required for myelin formation and maintenance is most vulnerable to alteration in β−tubulin 4 in those oligodendrocytes that form myelin on small diameter axons (Song et al., 1999).

### Preservation of axons in *taiep* dorsal columns

Despite the overt loss of myelin in the CST of 24-month-old *taiep* rats, we observed little evidence of axonal degeneration (pale, watery axoplasmic degeneration or dark dense axoplasm) or injury (accumulated organelles) (Dutta et al., 2006; Stassart et al., 2018; Schäffner et al., 2021). In the core of the CST and the F. gracilis, most axons appeared normal by TEM (**Fig. 3d and 4a**), with a pale axoplasm, neurofilaments and microtubules, and between 0 and 5 mitochondria per axonal cross-section. To examine axon viability across the *taiep* lifespan, we focussed again on the F. gracilis and the F. cuneatus and compared total axonal densities, axoplasm volume and axonal mitochondrial volume with WT.

**Fig. 4.**
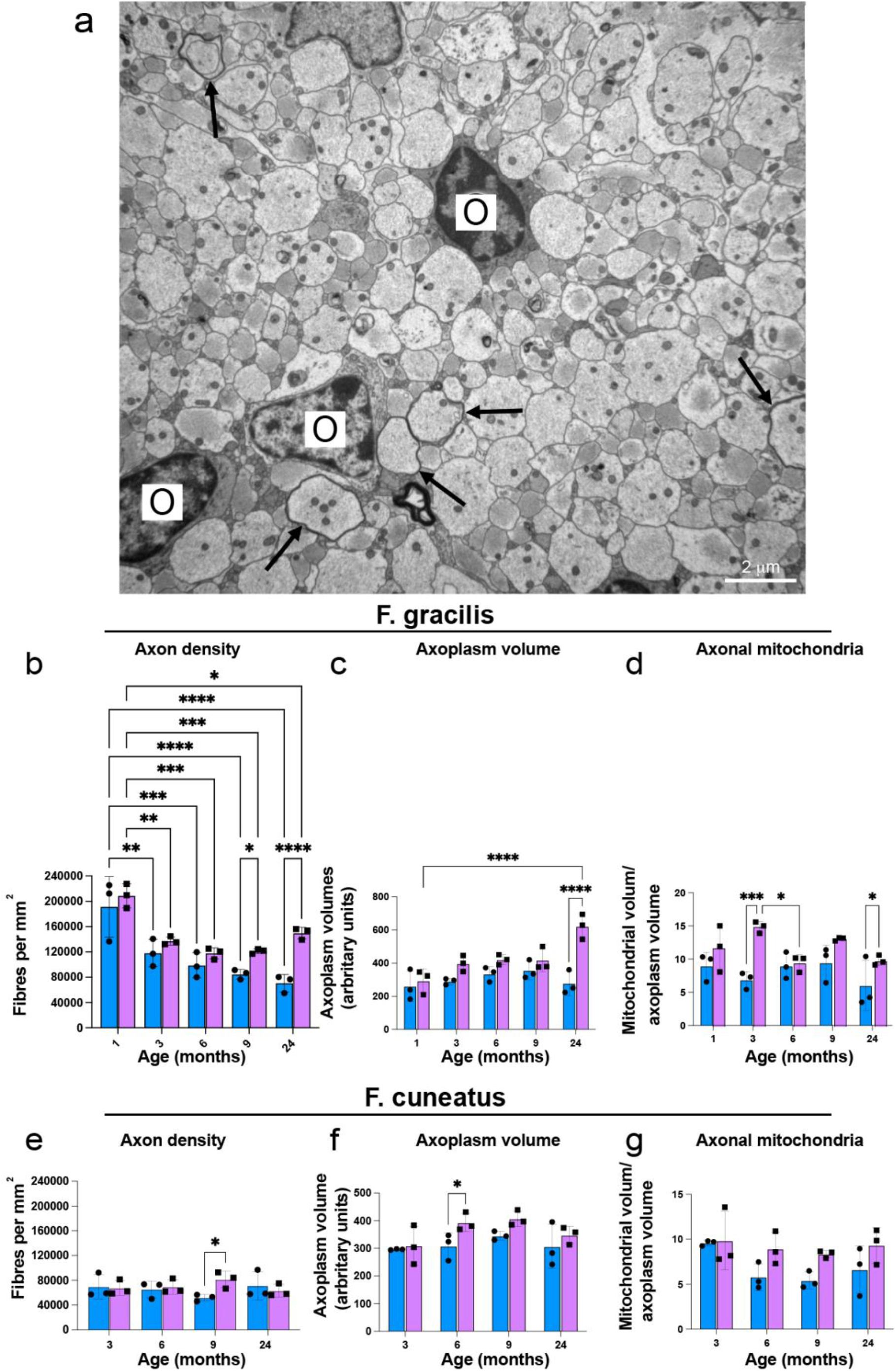
Preservation of axons in 24 month old *taiep* rat spinal cord dorsal columns. **a.** TEM micrograph of 24-month-old *taiep* F. gracilis. In the core of the F. gracilis, most axons of all calibers were demyelinated, but the density of axons appeared normal. A few axons however had thin myelin sheaths (black arrows). O: Oligodendrocyte nucleus. **b.** Graph of axonal densities in the F. gracilis at 1 month through 24 months in WT and *taiep*. Densities were similar in both genotypes at each timepoint except at 9 and 24 months when densities were significantly higher in *taiep* compared to WT, most likely due to the tight packing of demyelinated axons. A significant decrease in densities compared to 1-month-old animals was observed in both genotypes, as expected given the growth in fibre size with adulthood. **c.** Axoplasm volume tended to be higher in *taiep* compared to WT at all time points, reflecting the reduction in space occupied by myelin. Differences reached significance only at 24 months. In *taiep,* the axoplasm volume was significantly higher at 24 months compared to 1 month. Since axonal densities changed in the opposite direction, this likely reflects both increased axonal size and loss of space occupied by myelin **d.** Mitochondrial volumes tended to be higher in *taiep* axons than WT axons at all ages, reaching significance at 3 and 24 months. **e.** In the F. cuneatus, axon densities were similar in WT and *taiep* at all time points except at 9 months when densities in *taiep* were significantly higher than WT. This likely reflects a statistical anomaly given that WT densities appeared untypically low compared to other ages. **f.** Axoplasm volume tended to be higher in *taiep* compared to WT from 3 months of age, reaching significance only at 3 months. This likely reflects loss of space occupied by myelin. **g.** Axon mitochondrial volumes tended to be higher in *taiep* compared to WT at 6, 9 and 24 months, but the differences were not significant. All graphs: black dots represent individual animals and bars represent means +/- SD.

F. gracilis: Comparing WT and *taiep* to themselves over time, total axon densities were significantly decreased in both genotypes at 3, 6, 9 or 24 months compared to 1 month, reflecting the growth in the cross-sectional area of fibres in adults compared to juveniles (**Fig. 4b**). The axoplasm volume (reflecting the relative space occupied by axoplasm compared to other white matter elements) was also significantly increased in 24-month-old *taiep* rats compared to 1-month-old animals, likely reflecting a combination of axon growth and loss of space occupied by myelin (**Fig. 4c**). Comparing genotypes, the density of axons was significantly higher in *taiep* than WT at 9 and 24 months, likely due to tighter packing of demyelinated axons (**Fig. 1f**). Correspondingly, axoplasm volume was significantly higher in *taiep* than WT at 24 months (**Fig. 4c**). Mitochondrial volume (normalised to axoplasm volume) tended not change over time in either genotype, despite being significantly higher at 3 months than 6 months, in *taiep*. Comparing genotypes however, mitochondrial volume tended to be higher in *taiep* compared to WT at all time points, achieving significance at 3 months when demyelination begins, and at 24 months when most axons were demyelinated (**Fig. 4d**).

F. cuneatus: Comparing the density of axons in WT and *taiep* to themselves over time, we found no differences. Comparing genotypes, axonal densities were also similar at each time point, although at 9 months the density in *taiep* was slightly but significantly higher than WT (**Fig. 4e**). Correspondingly, axoplasm volume did not change over time in either genotype. Nonetheless, when comparing genotypes, axoplasm volume tended to be elevated in *taiep* compared to WT at 6, 9 and 24 months, reaching significance only at 6 months (**Fig. 4f**). Axonal mitochondrial volumes also did not change significantly with time or with genotype, despite demonstrating an upward trend in *taiep* compared to WT from 6 months (**Fig. 4g**).

### Supernumerary glial cell processes and oligodendroglia hyperplasia in *taiep* dorsal columns

A notable observation in 24-month-old *taiep* was the presence of supernumerary glial cell processes interspersed between the axons. The extent of the area occupied by these processes is illustrated by pseudocolouring in **Fig. 5a**. Astrocyte processes (green), identified by densely packed glial fibrils, occasionally appeared partially translucent and contained small dark puncta consistent with glycogen granules, a characteristic of this cell type (inset top right **Fig. 5a**). Oligodendrocyte processes (yellow), were identified by their microtubular content and absence of neurofilament. Additionally, there was a clear qualitative increase in oligodendrocytes in *taiep* (**Fig. 5b & c**) compared to WTs. In one-micron thick resin sections of 24-month-old *taiep* rats, the majority of oligodendrocytes had medium/large round nuclei with the appearance of Type 1/2 oligodendrocytes as defined previously (Mori and Leblond, 1970) (**Fig. 5b & c**). TEM confirmed these cells were oligodendrocyte as they were a uniform population with increased microtubule densities (**Fig. 5d**). Quantitation of oligodendrocytes, astrocytes and microglia (as defined in **Supplementary Fig. 5**) showed a major increase in oligodendrocyte densities in *taiep* compared to WTs (**Table 1**). Consistent with a gain of toxic function mutation in *Tubb4a*, accumulations of microtubules (**Fig. d-j**) were found throughout the oligodendrocyte cytoplasm, as parallel arrays extending from the soma into cell processes (**Fig. 5g**) and in both large and small processes. Almost all oligodendrocyte somata had arrays of smooth endoplasmic reticulum (SER) coated by microtubules, as reported previously (Couve et al., 1997) (Duncan et al., 2017). Microtubule attachment to the SER began as the SER arose by extension of the outer nuclear envelope (**Fig. 5h-j**).

**Fig. 5.**
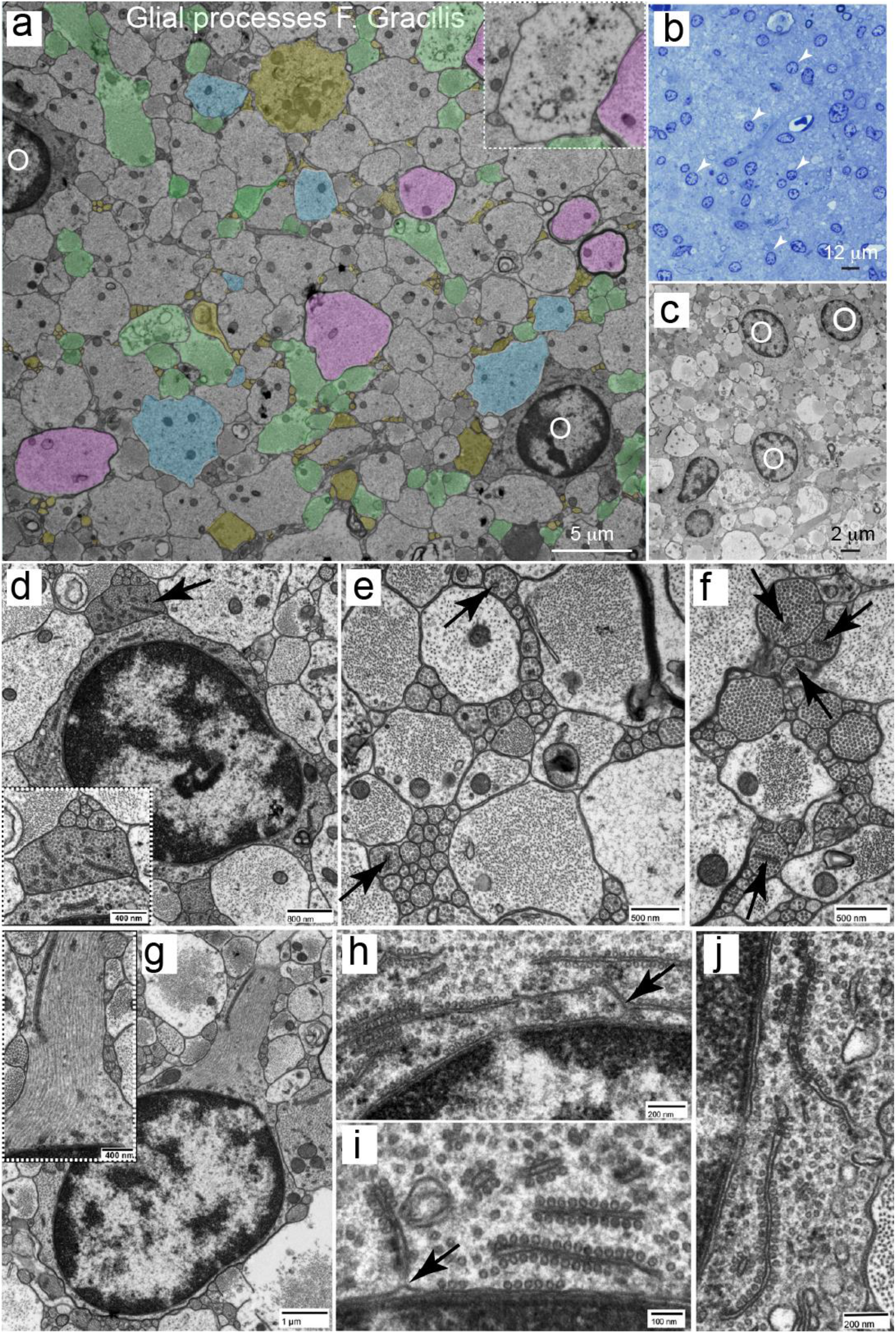
Supernumerary glia and glia cell processes in 24-month-old *taiep* rat. **a.** Electron micrograph of *taiep* F. gracilis at 24 months pseudo coloured to illustrate the extent of glial processes. Astrocyte processes (green) and oligodendrocyte processes (yellow) occupied a large amount of space between demyelinated (blue) and thinly myelinated axons (magenta). In the inset, putative glycogen granules can be observed in an astrocyte process. OL: oligodendrocyte soma; Ax: Axon; My Ax: Myelinated Axon; As: Astrocyte **b.** Light microscopy of the toluidine blue stained *taiep* F. gracilis shows many presumptive oligodendrocytes with round nuclei (arrows) are present in the demyelinated tract. Scale bar: 12 μm. **c.** On TEM, these cells can be seen to have accumulated microtubules, confirming that they are oligodendrocytes (O). Scale bar: 2 μm. **d.** All identified oligodendrocytes had accumulation of microtubules (MTs) throughout the cell cytoplasm and many MTs were aligned along cisterns of smooth endoplasmic reticulum (SER). Adjacent to such cells, oligodendrocyte processes (arrow, see inset) with MT accumulation, were frequently seen (inset). Scale bars: 800 nm, inset 400 nm. **e&f.** Formation of microprocesses. Collections of oligodendrocyte microprocesses of varying calibre were frequently noted, interdigitated between demyelinated axons and astrocyte processes. In the processes, MTs were frequently aligned along SER (arrows). Scale bars: 500 nm. **g.** Microprocesses arise from oligodendrocytes. Parallel arrays of tightly packed MTs were seen in large processes which led away from the oligodendrocyte cell body. Scale bars: 1um, inset 500 nm **h-j.** Distribution of MTs within the oligodendrocyte cell body. MTs frequently aligned along the outer nuclear envelope (arrowhead) and also along outpouches of the nuclear envelope which projected into the cytoplasm (arrows). These outpouches developed into free cytoplasmic cisternae that were decorated with MTs, sometimes with gaps along the SER surface (g). Scale bars: h 200 nm, i 100 nm, j 200nm.

**Table 1.** Cell densities in the F. gracilis at 24 months of age, identified by nuclear characteristics (Supplementary Fig. 5)

|  | WT | *Taiep* |
|---|---|---|
| Oligodendroglia | 10 (+/- 4.6 St dev; n = 3) | 67 (+/- 10.7 St dev; n = 3) |
| Astrocytes | 1.7 (+/- 2.1 St dev; n - 3) | 3.7 (+/- 3.1 St dev; ne = 3) |
| Microglia | 1.7 (+/- 0.6 St dev; n = 3) | 3 (+/- 2.0 St dev; n = 3) |

### Microglial activation in the *taiep* rat dorsal column accompanies myelin loss

Using the OX-42 antibody in spinal cord sections at 3, 6 and 13 months, there was a marked qualitative increase in microglial densities in *taiep* compared to WT (**Fig. 6a-f**). In *taiep* F. gracilis and CST, most appeared amoeboid in shape. In contrast, in the *taiep* F. cuneatus, microglia remained ramified, as in all the WT samples (**Fig. 6a-c**). Due to a lack of tissue from older rats, we were unable to examine the microglial density and morphology at 24 months, when most axons are demyelinated.

**Fig. 6.**
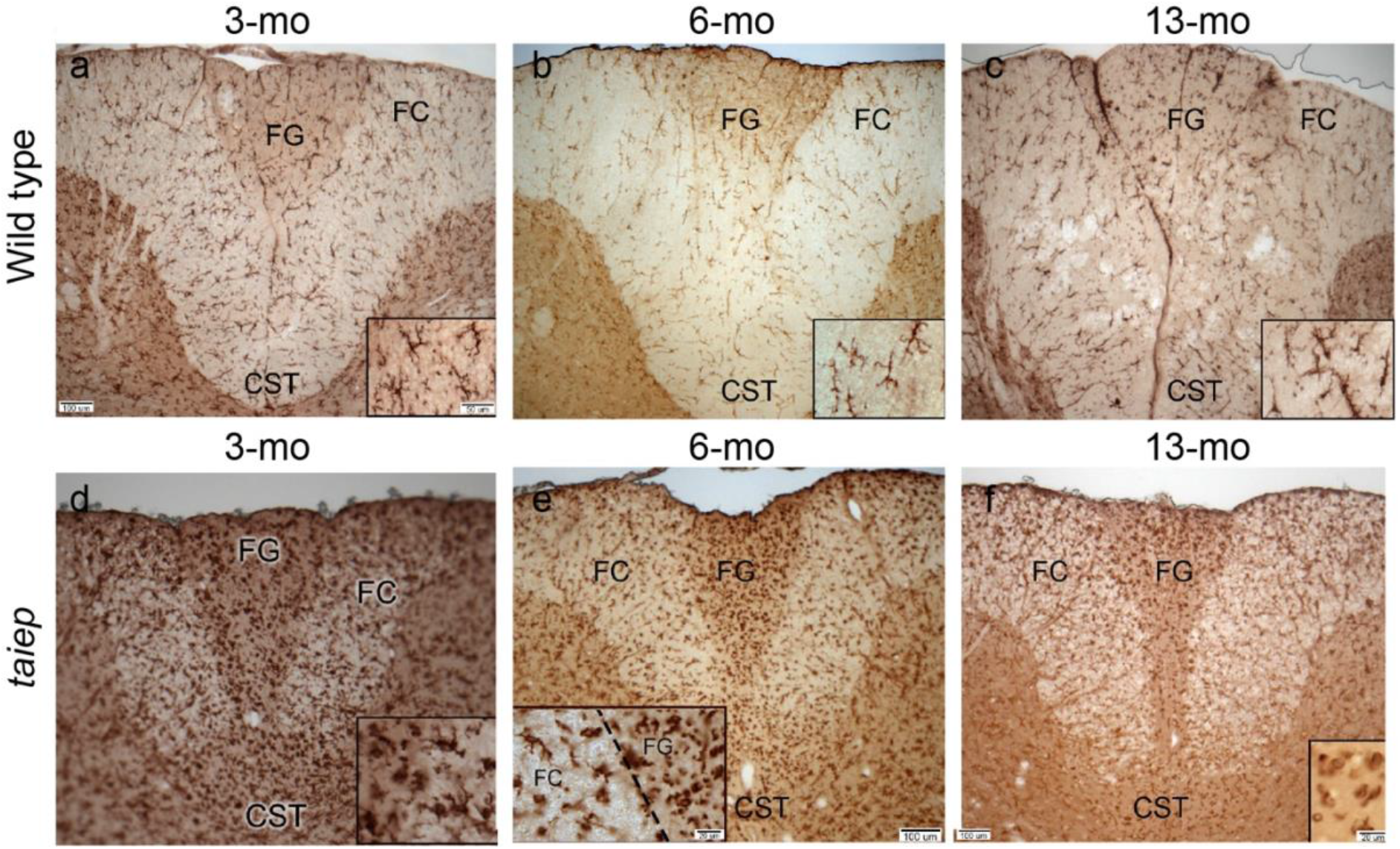
Microglial activation is most pronounced in tracts undergoing severe demyelinationa-c. At 3, 6 and 13 months of age in WT rats, microglia throughout the dorsal columns appear ramified indicating their resting state (CD11b immunolabelling). **d-f.** However, in the F. gracilis and CST of age-matched *taiep* rats, microglia appear increased in density and appear ‘activated’ as indicated by their amoeboid appearance. In the insets in E there is a clear demarcation between the F. gracilis (FG) and F. cuneatus (FC); in the latter microglia appear ramified.

## Discussion

The principal finding of this study is that small-diameter spinal cord axons can survive into old age despite prolonged and extensive demyelination. In the *taiep* rat, the progressive decline in densities of myelinated axons, beginning between 3 and 6 months of age, was accompanied by a corresponding increase in non-myelinated axon density, consistent with the loss of pre-existing myelin. By 24 months, only ∼10% of F. gracilis axons remained myelinated, yet total axon density was comparable to that of age-matched WTs. Importantly, axons appeared largely intact ultra-structurally, aside from the expected increase in mitochondrial volume. A near- complete loss of myelin was also observed in the CST and affected small diameter axons of the F. cuneatus. Collectively, these findings demonstrate that chronic demyelination does not inevitably result in axonal loss and suggest that myelin is not invariably required for long-term axonal survival.

These results extend our and others’ previous observations in MS, experimental autoimmune encephalomyelitis (Schäffner et al., 2021) and *Plp1*-mutant mice (Edgar et al., 2010; Groh et al., 2023) in which acute demyelination was compatible with axonal preservation. Here we substantially extend the period over which survival of demyelinated axons has been demonstrated, hitherto in Long-Evans shaker rat, a myelin basic protein gene (*Mbp*) mutant, in which demyelinated spinal cord ventral column axons survive until the rat is 9 months of age (Smith et al., 2013).

In MS, progression independent of relapse activity (PIRA) can continue throughout both the relapsing-remitting and progressive phases of the disease (Kappos et al., 2020; Lublin et al., 2022). PIRA is associated with increased spinal cord atrophy and a greater burden of paramagnetic rim lesions, both markers of neurodegeneration, highlighting the need for understanding the mechanisms underlying neuronal and axonal injury in MS. It has been proposed that the loss of chronically demyelinated axons in MS results from sustained energetic demands associated with continuous impulse conduction, often described as a state of "virtual hypoxia," together with the loss of oligodendroglial trophic and metabolic support (Nave and Trapp, 2008; Trapp and Nave, 2008; Trapp and Stys, 2009; Mahad et al., 2015).

The present findings suggest that these consequences are not inevitable, although an important limitation is that we did not determine whether the surviving dorsal column axons remained electrically active. Nevertheless, it is noteworthy that the distal segments of small-diameter axons within the cervical F. gracilis possess characteristics that make them particularly susceptible to disruptions in axonal transport and energy metabolism, as demonstrated in several experimental and pathological settings (Garbern et al., 2002; Edgar et al., 2004; Coleman, 2005; Bridge et al., 2009) including MS (DeLuca et al., 2004). Thus, their resistance to demyelination-induced degeneration in the *taiep* rat is particularly remarkable.

At first sight, our findings appear to differ from reports of substantial optic nerve axon loss in *taiep* rats at 12 months and following long-term toxin-induced demyelination in macaques (Wilkins et al., 2010; Sarrazin et al., 2022). Several factors may explain this difference. Axonal vulnerability may vary between anatomical pathways according to axon calibre, length, firing pattern and metabolic demand, as well as the local glial environment. However, given the known susceptibility of cervical cord dorsal column axons (Garbern et al., 2002; Edgar et al., 2004; Coleman, 2005; Bridge et al., 2009), we speculate rather that the discrepancy lies in the method used for enumerating axons. Chronically demyelinated or metabolically stressed axons may alter neurofilament phosphorylation, neurofilament organisation or antigen accessibility and could consequently become indiscernible immunohistochemically, without having degenerated. In contrast, TEM provides direct identification of small axonal profiles, although it too is subject to sampling limitations, and importantly, axonal densities, while indicative, should not be understood to indicate that axonal numbers are preserved.

An important question is how chronically demyelinated axons survive. Axonal mitochondrial volume was increased significantly in the *taiep* rat F. gracilis at 3 and 24 months, corresponding respectively to the onset of demyelination and its advanced stage. Increased mitochondrial content is a recognised response to dys- and demyelination and may compensate for the increased ATP demand created by redistribution of ion channels and continuous impulse conduction (Andrews et al., 2006; Ohno et al., 2014; Licht-Mayer et al., 2020, 2023). Indeed, increased mobilisation of mitochondria from the neuronal cell body to the axon has been shown to be neuroprotective for vulnerable, acutely demyelinated axons (Licht-Mayer et al., 2020). The mitochondrial response we observed therefore confirms that the demyelinated axons were metabolically altered and should not be regarded as unaffected. Nevertheless, the scarcity of signs of axonal pathology such as axonal organelle accumulation (Stassart et al., 2018) or axoplasmic disintegration (Dutta et al., 2006) suggests that this compensatory response contributed to survival. Whether these axons continued to conduct impulses (even if slowly), or were electrically silent but structurally preserved, remains undetermined. Importantly, the energy demands of firing and non-firing axons are substantially different (Trevisiol et al., 2017; Looser et al., 2022) and axonal preservation alone is unlikely to be highly energy dependent.

The marked glial response may provide an additional explanation for axonal persistence. Oligodendrocyte density in the aged *taiep* F. gracilis was substantially greater than in WTs, and numerous oligodendrocyte and astrocyte processes were interwoven with demyelinated axons. Oligodendrocytes and astrocytes support axons through mechanisms that are separable from myelin formation, including the provision of trophic and/or metabolic substrates (Griffiths et al., 1998; Fünfschilling et al., 2012; Lee et al., 2012; Fulmer et al., 2014; Chamberlain et al., 2021; Duncan et al., 2021). We therefore hypothesise that persistent glial processes can transfer sufficient metabolic and/or trophic factors to sustain axons in the absence of myelin sheaths.

This interpretation is consistent with the accepted view that the oligodendrocyte–axon unit has functions extending beyond saltatory conduction. The accumulated oligodendrocyte processes likely represent a pathological consequence of the mutation rather than an adaptive response. Nonetheless, these possibilities are not mutually exclusive: morphologically abnormal oligodendrocyte processes might retain sufficient metabolic function to support axonal survival even though microtubule-dependent intracellular trafficking (Song et al., 1999, 2001, 2003) and myelin maintenance are impaired. Astrocyte processes, some containing puncta consistent with glycogen granules, could offer a source of energy substrates (Pellerin et al., 1998; Veloz Castillo et al., 2021), either directly or through oligodendroglial networks (Späte et al., 2024). Nonetheless, morphology alone cannot establish a neuroprotective function, and this possibility will require the demonstration of the expression and localisation of relevant metabolic transporters on these glial cell processes.

The presence of activated microglia is important when considering the relevance of this model to MS. Amoeboid-appearing microglia accompanied dysmyelination and ongoing demyelination in the F. gracilis and CST of *taiep* rat, where demyelination was most marked. Nonetheless, at 24 months, microglial densities in *taiep* F. gracilis were similar to controls suggesting substantially reduced neuroinflammation following clearance of myelin debris, as we described previously in *Plp1* transgenic mice [20]. In a murine myelinating cell culture system, myelin debris clearance by microglia is associated with increased levels of C-C Motif Chemokine Ligand (CCL) 3 and 4 (Bijland et al., 2019) and a recent study by the Simons lab (Berghoff et al., 2026) showed that chronic myelin pathology induces progressive accumulation of a pro-inflammatory, myelin-damaging microglial state marked by increased *Ccl3/Ccl4* expression and impaired cholesterol processing. Assuming a similar response in actively demyelinating *taiep* rat F. gracilis and CST, this did not reach a threshold sufficient to cause widespread injury or degeneration of axons themselves (Smith et al., 2001; Nikić et al., 2011).

The vulnerability of *taiep* rat myelin showed a striking relationship to axon calibre; small-diameter axons being severely affected, while many large-diameter axons retained apparently normal myelin even at 24 months. This pattern is unlikely to be explained simply by the absence of ‘toxic’ β-tubulin 4 from those oligodendrocytes myelinating large diameter axons. Rather, our earlier studies suggest that small axons, myelinated by Type1/2 oligodendrocytes that have multiple fine processes are at greatest risk for demyelination. We previously showed that, 1) the polarity of microtubules in fine-calibre oligodendrocyte processes is altered (Song et al., 1999) 2) there is a trafficking defect in *Mbp* mRNA to the periphery in cultured *taiep* oligodendrocytes (Song et al., 2003), 3) in the anterior medullary velum (AMV), where myelination by either Type1/2 or 3/4 oligodendrocytes can be discerned, the part of the AMV myelinated by the fine, numerous processes of Type 1/2 cells, is preferentially disrupted (Song et al., 2001). Taken together, these data indicate that axonal size and their associated oligodendrocyte sub-type, are the key determinant that dictate the susceptibility to demyelination in the *taiep* rat.

One crucial limitation of our study must be considered. Structural preservation of axons does not establish normal conduction or neurological function: *taiep* rats remained severely neurologically impaired despite the survival of many axons. This qualification is particularly relevant for extrapolation to MS. Human MS lesions are heterogeneous and expose axons to varying combinations of inflammation, oxidative injury, mitochondrial dysfunction and loss of glial support. The present findings do not imply that chronic demyelination is benign, nor do they diminish the importance of remyelination (Franklin and Ffrench-Constant, 2008; Duncan et al., 2024). Instead, the results suggest that there may be a prolonged interval during which chronically demyelinated axons remain structurally viable and potentially amenable to remyelination (Duncan et al., 2009). Therapies that preserve or enhance the putative non-myelinating supportive functions of oligodendrocytes and astrocytes might extend this therapeutic window, particularly where remyelination is delayed or incomplete.

Our findings are compatible with two scenarios: first, that demyelinated axons lose their dependence on myelin-mediated trophic and metabolic support and can survive in its absence possibly taking up energy substrates directly from the extracellular milieu, or second, that glia, in intimate contact with demyelinated axons, can exert such support though monocarboxylate transporters on glia and axons. Compact myelin remains essential for efficient conduction and likely for optimal axonal protection, but chronic myelin loss alone is not necessarily sufficient to cause axonal death.

## Contributions

Devised the study and managed all aspects: IDD

Drafted the manuscript, analysed and interpreted the data: JME; NL.

Contributed text: IDD

Conducted experiments: IDD; BA; NL; DR, KJC; HA; JME

Generated images: IDD, BA and JME

Figures: JME

Edited and approved the manuscript: all authors

## Supplementary Information

**Supplementary Fig. 1.**
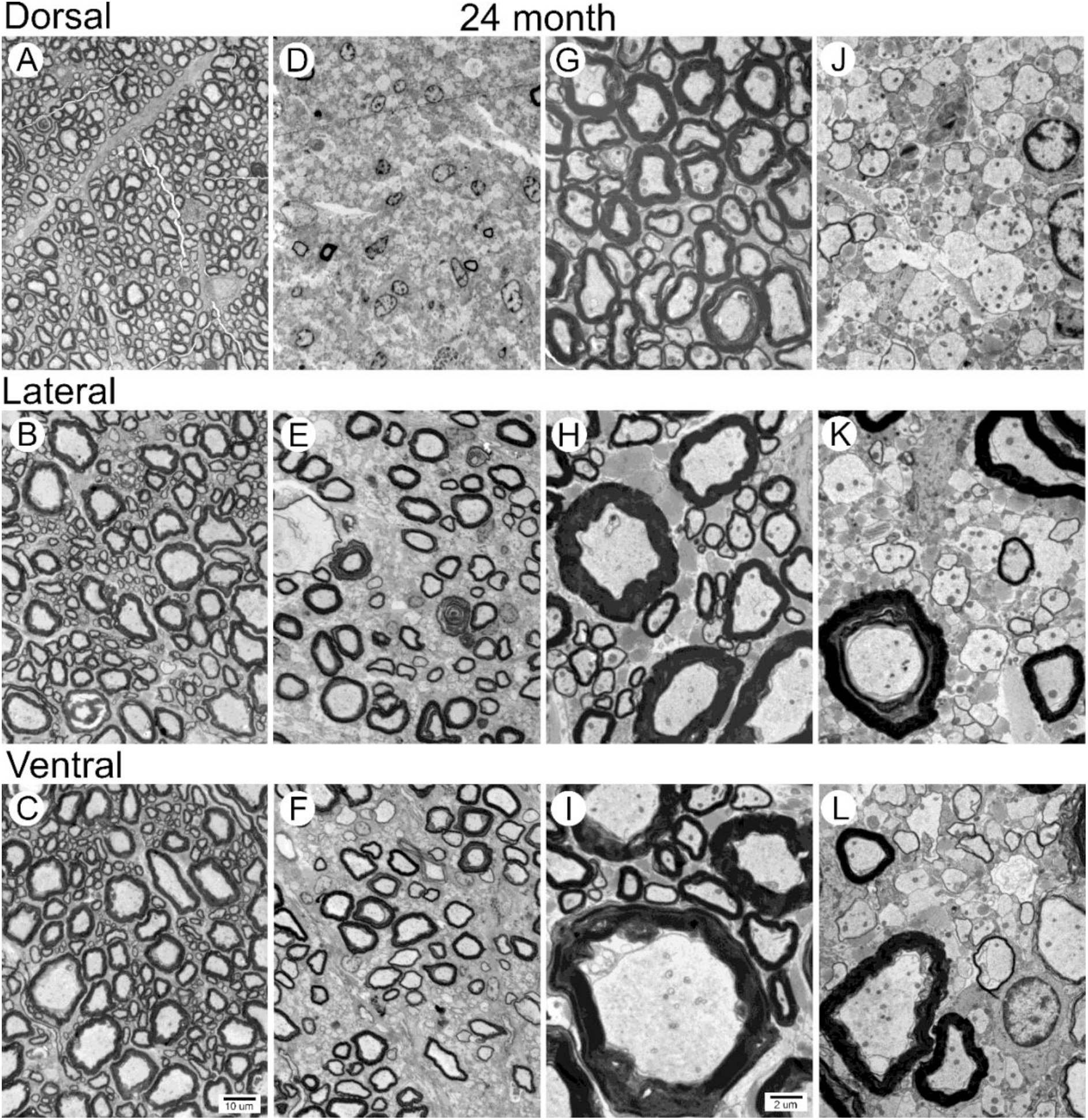
Electron micrographs of Dorsal, Lateral and Ventral columns in the cervical cord of WT and taiep rats at 24 months of age, demonstrating the vulnerability of small diameter axons to demyelination.

**Supplementary Fig. 5.**
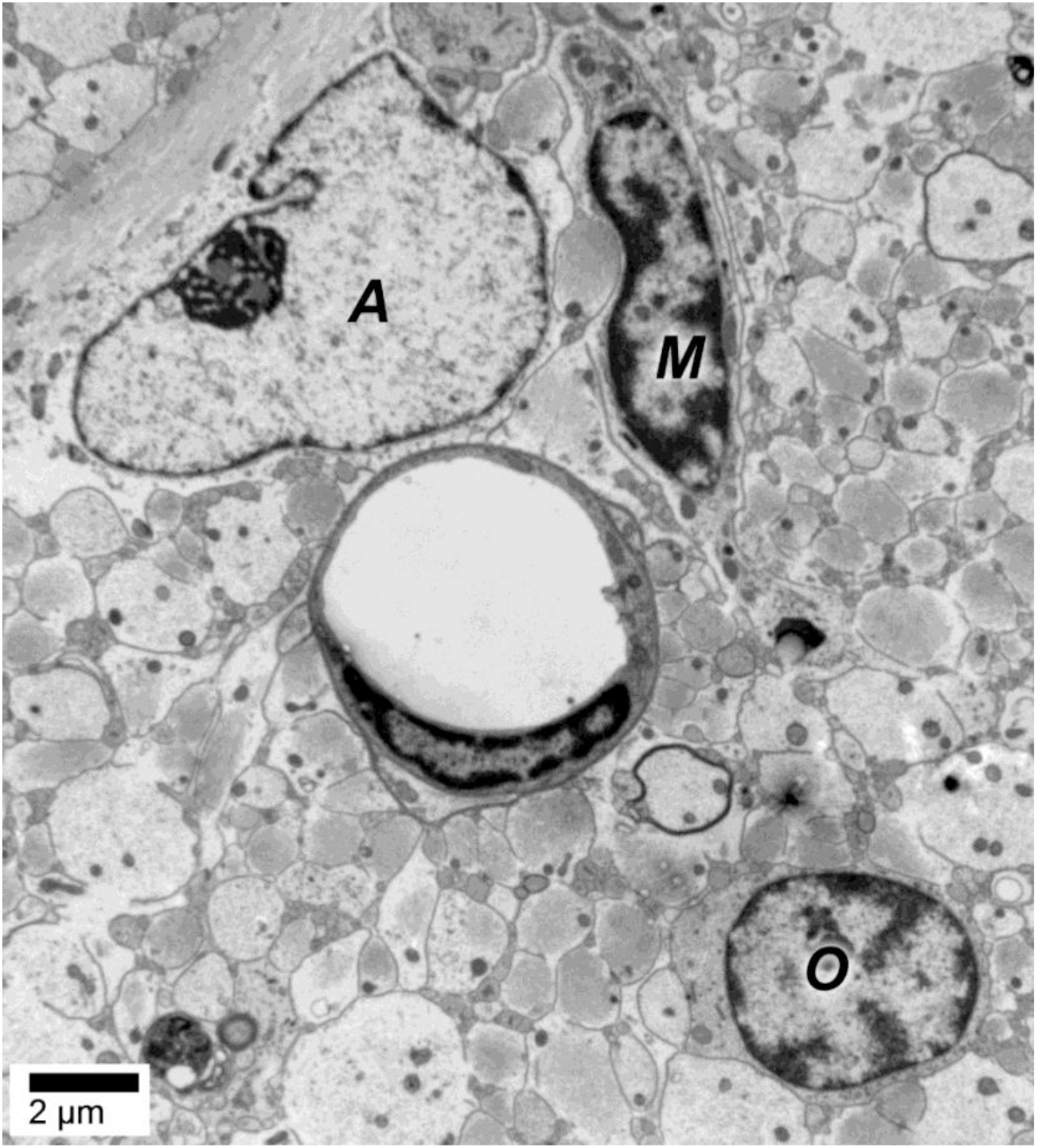
There is a clear and unequivocal difference between oligodendrocytes (O) with dense chromatin in their nuclei and, in *taiep*, microtubules in the perinuclear cytoplasm, astrocytes (A) with pale nuclei with little dense chromatin and microglia (M) which tend to have bean-shaped nuclei, to allows their ultrastructural quantification. The hollow lumen in the middle belongs to a blood vessel. Scale bar: 2 μm.

